# Comprehensive-immunopeptidomics analysis reveals presentation of potential neoantigens carrying cancer driver mutations

**DOI:** 10.1101/2021.04.09.439118

**Authors:** Yuriko Minegishi, Kazuma Kiyotani, Kensaku Nemoto, Yoshikage Inoue, Yoshimi Haga, Risa Fujii, Naomi Saichi, Satoshi Nagayama, Koji Ueda

## Abstract

Understanding the properties of human leukocyte antigen (HLA) peptides (HLAp) is essential for cancer precision medicine, while the direct identification of HLAp from a tiny piece of clinical tissues by mass spectrometry (MS) is still confronted with technical challenges. Here, to overcome that hindrances, high-field asymmetric waveform ion mobility spectrometry (FAIMS) is introduced to conduct differential ion mobility (DIM)-MS by seamless gas-phase fractionation optimal for scarce samples. The global-immunopeptidomics analysis enables in-depth identification of 44,785 HLAp, including 2 neoantigens with KRAS-G12V and CPPED1-R228Q from stage IV colorectal cancer (CRC) tissues obtained from 17 subjects. Comparison of tissue-based personal immunopeptidome disclosed cancer-specific processing of HLAp. Since the direct identification of neoantigens from CRC tissues suggests more potential neoantigens have yet to been identified, the targeted-immunopeptidomics by screening-oriented parallel reaction monitoring (PRM) has been established and identified two more neoantigens with oncogenic KRAS. The targeted-immunopeptidomics analysis clarifies the presentation of aimed neoantigens, including neoantigens that are unpredictable by computational algorithms. The comprehensive-immunopeptidomics approach combining the FAIMS-assisted DIM-MS and the screening-oriented PRM contributes the direct and effective identification of HLAp and emerging neoantigens from clinical tissue samples, as an advanced strategy to identify the authentic and recurrent neoantigens for cancer immunotherapy.

## 1. Introduction

Human leukocyte antigen (HLA) is the major histocompatibility complex (MHC) of human that plays a role in self-tolerance and innate immunity. The HLA complex contains an HLA peptide (HLAp, also called immunopeptide) that plays a role in self-nonself discrimination. Neoantigens are one of the HLAp that carry cancer-specific somatic mutations. The neoantigens presented on the surface of cancer cells are recognized as nonself antigen by the relevant repertoire of T cells and can trigger anti-cancer immunity.^[1]^ The immune checkpoint inhibitors (ICIs) have been established to invigorate this anti-cancer immune system as cancer immunotherapies.^[2]^ Although immunotherapies are efficacious against certain cancer types,^[2]^ there is still a lack of knowledge about predictive factors for successful ICI indication.^[2–3]^ From these perspectives, the analysis of pathologically-independent tissue-based immunopeptidome from identical patient is increasingly important to predict the efficacy of ICI in patients, further explore the new targets for therapeutic intervention.

Mass spectrometry (MS) immunopeptidomics is currently the only method that can directly detect HLAp. According to the guidelines for immunopeptidomics,^[4–5]^ one of the disadvantages of MS-based immunopeptidomics is incapability of robust analysis from a scarce sample, such as tiny tissue of endoscopic biopsy, compared to the genomic prediction of HLAp by *in-silico*.^[6–8]^ In the previous method for immunopeptidomics, in order to obtain thousands of convincing immunopeptidome, more than 1e^8^ of cultured cells or nearly 1 gram of tissue samples were required.^[6–8]^ In addition, to ensure the best results, the process of chemical pre-fractionation is inevitable, that will become an obstacle to the analysis from scarce samples. In fact, direct detection of neoantigens from solid tumors (such as colorectal, liver, or ovarian cancer) has been unsuccessful before.^[5, 9–10]^ Due to the cancer-associated microenvironment, the diversity of HLAp at the tissue level can be dynamic. Therefore, understanding the cancer immunopeptidome at the tissue level with the appropriate pathological control from the same individual is essential. Establishing the more efficient approach that enables the in-depth immunopeptidomics analysis from scarce tissues is dispensable for the field of advanced precision cancer immunotherapy.

It has been suggested that neoantigens will be depleted during tumor development.^[11–14]^ This is because the immunogenic neoantigen-presenting cells are eradicated by the anti-cancer immune system and are thought to hardly remain in advanced cancers. Especially, since the cancer driver mutation is necessary for the onset of carcinogenesis from the very early stage, the cancer cells that present cancer driver mutation-carrying neoantigens are thought to be depleted in the cancer progression to the late stage. Therefore, neoantigens carrying cancer driver mutations are rarely exist in advanced cancer tissues and are rarely directly identified by mass spectrometry. The low abundance of neoantigens in cancer tissues may make it difficult for MS to directly identify neoantigens from advanced cancer tissues. In contrast, a recent report pointed out the lack of necessary normalization in the previous genomics analyses of neoantigen depletion, and that led to a false-positive signal of neoantigen depletion. This pseudo-signal of neoantigen depletion was indeed cancelled when appropriate normalization was applied into the calculation.^[15]^ Therefore, the existence of neoantigens in advanced cancers, especially neoantigens with cancer driver mutations, is still controversial.

In this study, in order to solve the current task of immunopeptidomics, we adopted the differential ion mobility (DIM) -MS equipped with the interface of high-field symmetric waveform ion mobility mass spectrometry (FAIMS), suitable for the analysis of scarce samples. The seamless gas phase fractionation by FAIMS interface ensured a sufficiently deep immunopeptidome, so that for the first time, neoantigens became detectable directly from colorectal cancer (CRC) tissue samples. Since one of the neoantigens identified from CRC tissue carried oncogenic KRAS (G12V), a major cancer driver mutation, we further established the screening-oriented targeted-immunopeptidomics to elucidate whether potential neoantigens carrying oncogenic KRAS mutation are presented in cancer cells. As a result, two neoantigens carrying KRAS-G12V and one neoantigen carrying KRAS-G13D were identified by targeted-immunopeptidomics approach. Here, we show how comprehensive immunopeptidomics can help provide new insights into the pathological specific processing of immunopeptides and the presentation of potential neoantigens.

## 2. Results

### 2.1. Efficient identification of HLAp from scarce samples by newly established global-immunopeptidomics analysis

First, we fully optimized the parameters for DIM-MS to maximize the identification number of HLAp by the setting of three-fractionations at a time. In particular, a set of 9 compensation voltages (CVs), from −80 to −35 V, were selected and combined for the best performance of gas phase fractionation of HLAp (**Figure 1A**). The FAIMS-assisted immunopeptidome analysis using 5e^6^ cells of HCT116 identified an average of 3,215 HLAp. Resulting in a 1.3-fold increase over the condition without FAIMS (average or 2,482 HLAp, **Figure 1B**). Accordingly to the increase of HLAp, the number of identified source proteins was also greater in the FAIMS-assisted condition (2,381 proteins on average) than the without-FAIMS condition (2,077 proteins on average, Figure 1B). This result was consistent throughout the experiments and became statistically significant when the sample scale swas increased to 1e^7^ cells (average 3,893 HLAp identifications) or 1.5e^7^ cells (average 4,160 HLAp identifications, Figure 1B). By expanding the sample size under FAIMS conditions, the frequency of neoantigen detection was correspondingly increased up to 2.4 times (**Figure 1C**). Thus, gas-phase fractionation using DIM-MS can enhance the identification of HLAp and neoantigens, and even for samples that are an order of magnitude smaller, it can achieve efficient immunopeptidomics than most previous reports.^[6–7,9]^

**Figure 1.**
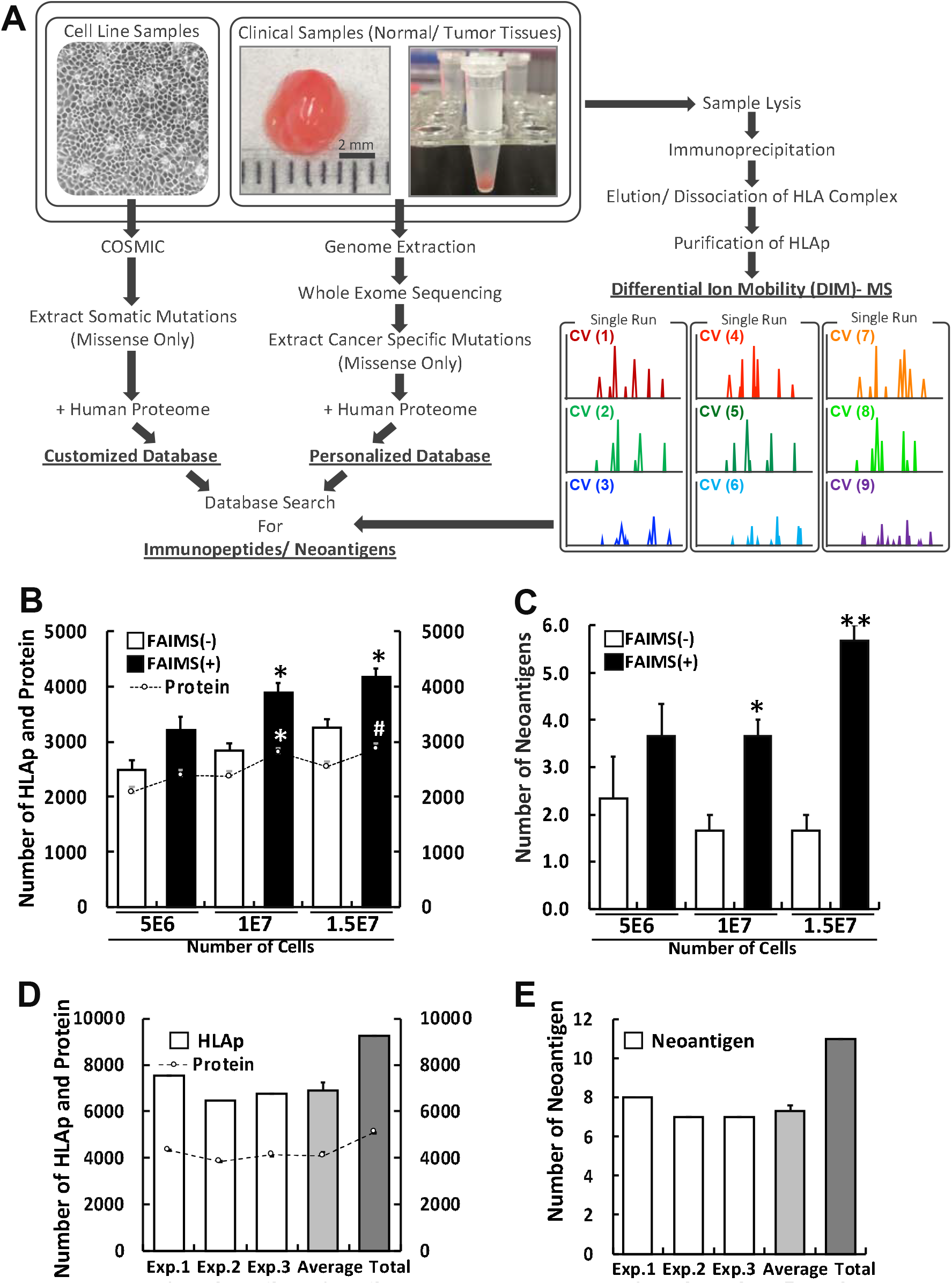
Schematic workflow and validation of global-immunopeptidomics analysis by FAIMS-assisted DIM-MS. A) The workflow for proteogenomic personalized immunopeptidomics analysis is shown. The DIM-MS analysis was performed under seamless gas-phase fractionations by installing the FAIMS-Pro interface. Acquired raw files were searched against the tailored database that includes normal human proteome with sample-corresponding cancer-specific somatic mutations. CV; compensation voltage. (B) The composite graph depicts the number of identified HLAp and the source protein in HCT116 cells. The horizontal axis represents the approximate number of cells used in each analysis. The bar graph and the vertical axis on the left (white bars; without FAIMS, black bars; with FAIMS) represent the number of HLAp. The dashed-line graph and the vertical axis on the right represent the number of identified source proteins from the identical analysis. * p < 0.05, # p < 0.1. (C) The average number of neoantigens identified from the identical experiments shown in (B) * p < 0.05, ** p < 0.01. (D) The composite graph depicts the number of HLAp identified in HCT116 by global-immunopeptidomics analysis. The number of HLAp identified from three independent analyses using 1e8 of HCT116 cells (white bars), the average number of HLAp per analysis (light gray bar) and the total number of unique HLAp (dark gray bar) are shown. The horizontal axis represents the three independent analyses, the average, and the total, respectively. The bar graph and the vertical axis on the left represent the number of HLAp. The dashed-line graph and the vertical axis on the right represent the number of identified source proteins. (E) The number of neoantigens identified from the identical experiments shown in (D).

From three independent analyses using 1e^8^ cells of HCT116, an average of 6,915 HLAp from 4,109 source proteins were identified, and a total of 9,249 unique HLAp were derived from 5,103 unique source proteins (**Figure 1D**). The length of and the restraint of identified HLAp exhibited the typical distribution and the assignment in the context of HLA allotype respectively (**Figure S1**). From this immunopeptidome, an average of 7.3 neoantigens were identified in each independent experiment, resulting in a total 11 of neoantigens (**Figure 1E**). Among these, 7 were not found in the immune epitope database (IEDB) (http://www.iedb.org/home_v3.php, as of November 27, 2020), hence regarded as newly identified ones (**Table 1**). These results indicated that the FAIMS-applied DIM-MS system could provide an ideal technological platform for in-depth profiling of HLAp, as well as direct detection of neoantigens.

**Table 1.**
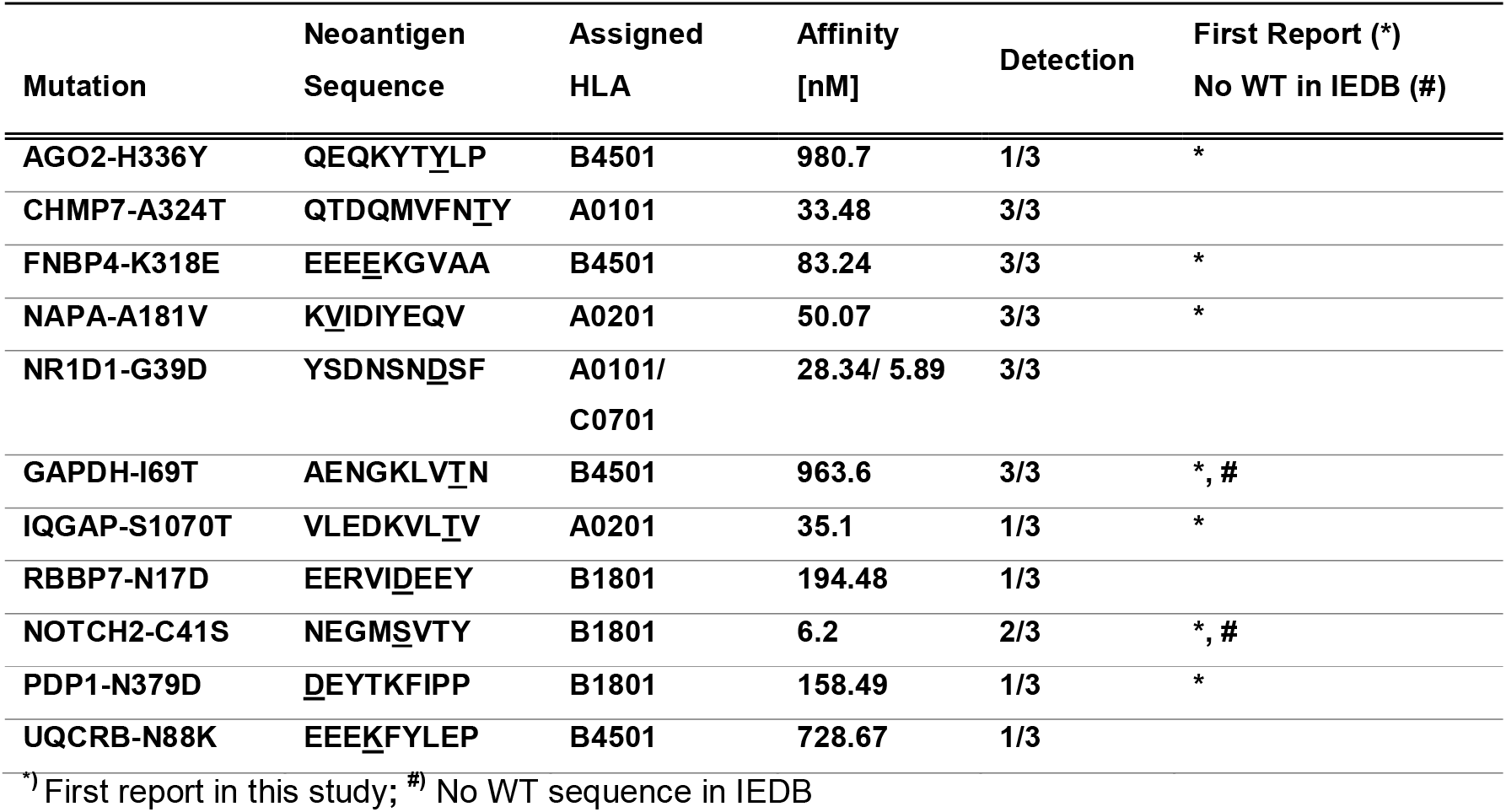
Neoantigens identified from HCT116 by global-immunopeptidomics.

### 2.2. Global-immunopeptidomics enabled direct identification of neoantigens from scarce tissues of CRC

By established global-immunopeptidomics, CRC samples of tumor or adjacent normal regions of CRC tissues from same individuals (n = 17) were subjected to personalized immunopeptidome analysis. By the analyses, a total, 44,785 unique HLAp were identified (6,578 peptides per patient on average) (**Figure 2A**). Among these, 5,603 HLAp (12.5%) were normal-exclusively identified and 14,052 HLAp (31.4%) were tumor-exclusive, while 25,130 HLAp (56.1%) were shared both in normal and tumor tissues. The comparison of immunopeptides in normal and cancer per patient can be found **Figure S2**. These results indicated that compare with the previous reports on the analysis of solid tumor tissues such as colon,^[5]^ liver,^[10]^ and ovarian cancer,^[9]^ the number of HLAp has been significantly improved when considering the required amount of sample here is smaller. From this CRC immunopeptidome, among 11,117 source proteins (average 4,149 proteins per patient), 817 (9.7 %) or 1839 proteins (16.5 %) were identified as normal or tumor-specific (**Figure 2B**). The number of immunopeptide identified from cancer tissues was significantly increased compare to the number of normal tissues, while it became insignificant when normalized by the amount of protein used for sample preparation (**Figure S3**). The identified HLAp exhibited the reasonable characteristics of HLA allotype-matched distribution and restraint by unsupervised clustering and the binding prediction (**Figure S4**). The length of HLAp was variable according to the HLA allotype in each individual, still, the typical dominance of Class I HLAp, 9-mer in length, was clearly shown and more than 99.7 % of immunopeptides were between 8 to 12 amino acids in length (**Figure S5**). Notably, 2 neoantigens were identified from 2 distinct patients’ tumor tissues (**Figure 2C**). The first neoantigen identified from the primary tumor tissue of ID 172 carried a well-known CRC driver mutation, KRAS-G12V, the position of the 7^th^ to 16^th^ amino acids (hereinafter, the position of amino acid will be described as [7-16] in square brackets). The other, serine/threonine-protein phosphatase CPPED1 has the R228Q mutation (CPPED-R228Q [226-234]), was identified from ID 261’s hepatic metastasized tumor tissue. KRAS-G12V is one of the representative cancer driver mutations, which can induce cancer in various tissues. While the other one, CPPED1, is known as a negative regulator of Akt signaling, which directly dephosphorylates Akt at Ser473.^[16–18]^ Since the amino acid substitution R228Q was found in the phosphatase domain of CPPED1, this mutation may attenuate the phosphatase activity and induce aberrant activation of Akt signaling in CRC cells. According to the motif information at IEDB and SYFPEITYHI (http://www.syfpeithi.de/0-Home.htm), the neoantigen carrying KRAS-G12V was predicted to have a higher affinity against A*11:01 than wild type sequence due to the preferred amino acid substitution of valine at position 6 in the epitope (Figure 2C, depicted as “Preferred: V” by an arrow). On the other hand, the mutation CPPED1-R228Q lost its deleterious amino acid of arginine at position 3 by substitution to glutamine (Figure 2C, depicted as “Deleterious: P/R” by an arrow). Therefore, compared with the corresponding wild-type (WT) sequence, the identified neoantigens respectively showed two opposite patterns, that is, by the acquired preferred residue, or the loss of deleterious residue in the epitope to obtain the affinity to A*11:01. This alteration reflected in the enhanced of affinity of in KRAS-G12V [7-16] than KRAS-WT [7-16] and CPPED1-R228Q [226-234] than CPPED1-WT [226-234] predicted as 137.3 nM vs 299.7 nM and 52.2 nM vs 729.9 nM, respectively.

**Figure 2.**
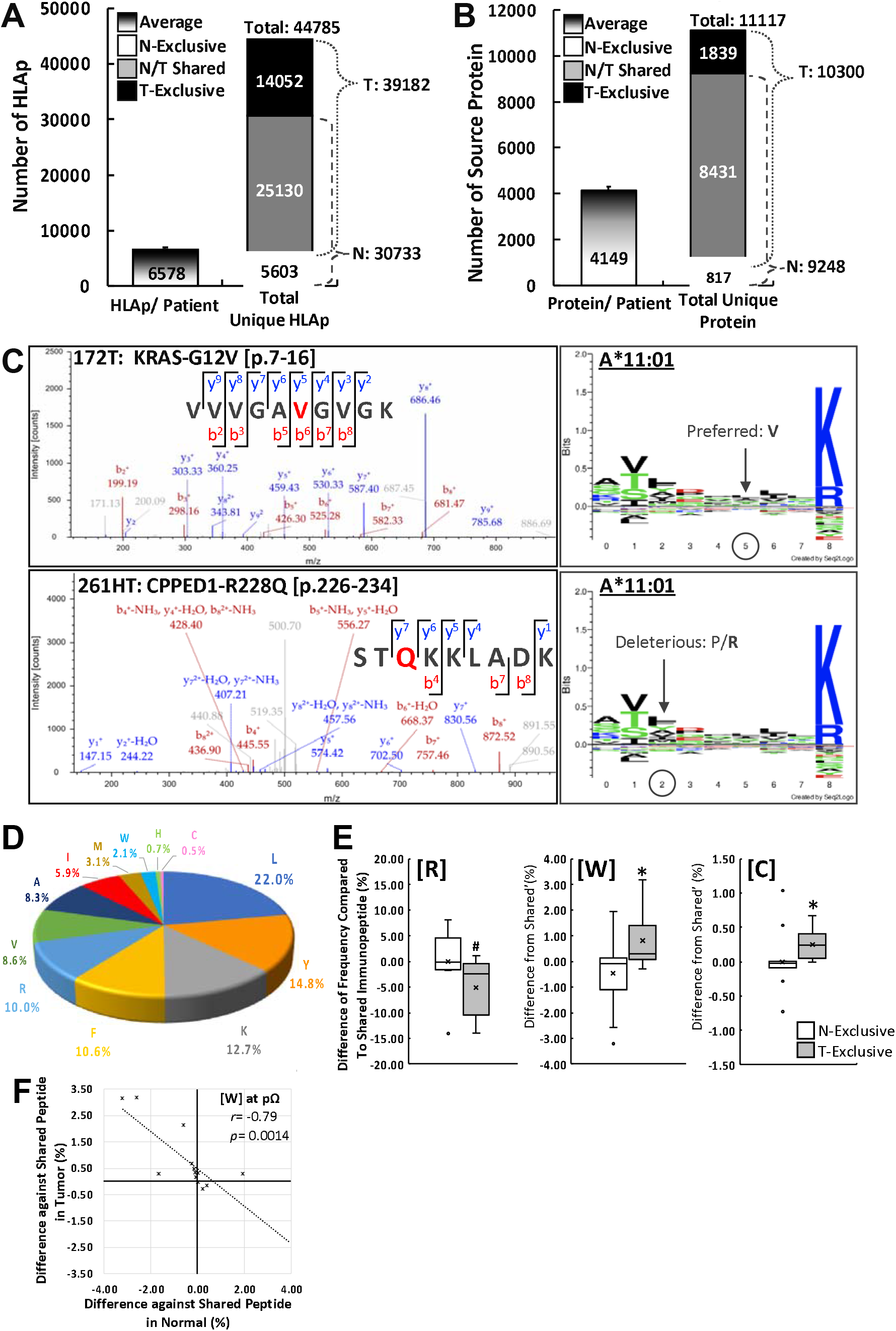
The CRC immunopeptidome, neoantigens directly identified from tissues and the cancer-specific profiling of HLAp by global immunopeptidomics analysis. A) A global-immunopeptidomics analysis was performed on the 17 sample sets (normal and tumor tissues) of independent patients with stage-IV CRC. The graph depicts the average number of HLAp identified from the normal and tumor samples per patient (HLAp/ Patient) and the total number of unique HLAp identified from 17 patients, *i.e.*, the immunopeptidome of CRC (Total Unique HLAp). For the profiling of cancer-specific HLAp, the CRC immunopeptidome was classified into three groups, namely, normal-exclusive (N-Exclusive), shared in both normal and tumor (N/T Shared), and tumor-exclusive (T-Exclusive). The numerals indicate the number of HLAp in each group. B) The graph depicts the average of the total number of source proteins per patient (Protein/ Patient) and the total number of source proteins from all patients (Total Unique Protein), identified by the same sample sets shown in A). C) The MS/MS spectra of neoantigens, KRAS-G12V [7-16] and CPPED1-R228Q [226-234], identified from CRC tissues. The substituted amino acid within neoantigen sequence is shown in red. The NetMHCpan4.1 predicted that these two neoantigens are the possible-binders of A*11:01. In the box on the right, the amino acid preference in the A*11:01 binding motif and the probable amino acid residues that affect the affinity for A*11:01 in each identified neoantigens are depicted. D) The pie chart depicts the frequency of amino acid usage at the C-terminus (pΩ) of HLAp. E) The box plot depicts the difference of frequency in the amino acid usage (%) of arginine (R), tryptophan (W) and cysteine (C) at pΩ of HLAp between the normal-exclusive and the tumor-exclusive groups compared to the shared-HLAp group. F) The negative correlation between normal-exclusive HLAp and the tumor-exclusive HLAp in the usage of W at pΩ of HLAp is depicted in box plot. The correlation was calculated by Pearson correlation coefficient.

### 2.3. Comparison of pathologically-exclusive immunopeptides revealed the cancer-specific profile of tryptophan trimming at pΩ

We next compared HLAp of three groups; HLAp only found in normal tissue (normal-exclusive), HLAp found in both normal and tumor tissues (shared), and HLAp only found in tumor tissue (tumor-exclusive) per patient. If there is a deviation in the antigen processing/presentation mechanism between normal tissue and tumor tissues, the distribution of HLAp should be shifted from the normal control. When we examined the usage for the C-terminus amino acid (pΩ) trimming for all the major population of immunopeptidome (8 to 12 amino acids in length), cysteine trimming exhibited the lowest frequency (0.5%) (**Figure 2D**). In contrast, tryptic (R, K) and chymotryptic (L, I, V, F, Y, W, A, M) amino acids were more common in the trimming of pΩ. Then, the usage of amino acid frequency at pΩ was calculated as ratio (%) against a subtotal of each group (normal-exclusive, shared, and tumor-exclusive peptide, respectively) and compared the difference against based on the frequency of the shared-peptide to clarify whether there would be a processing bias shown or not between normal-exclusive and tumor-exclusive groups. Among the 12 pΩ amino acids used, 9 patients carried at least one R-restraint HLA allotype (A*11:01 or A*33:03) at pΩ, and the usage of R in pΩ exhibited a tendency of reduction in tumor-exclusive HLAp (normal-exclusive vs tumor-exclusive: −0.06% vs −5.10% compared to the shared-peptide frequency, *p* = 0.0975, **Figure 2E**, [R]). Compare with the frequency of shared peptide, the usage of K exhibited no statistical difference (data not shown). Although R and K are in the same tryptic group, these two amino acids exhibited different dynamics in normal and tumor tissues. For chymotryptic peptides, unexpectedly, only W exhibited a statistically significant increase in tumor-exclusive HLAp, (Figure 2E, [W]). The pΩ trimming of HLAp by L, I, and V (usually classified as similar hydrophobic amino acids) exhibited inconsistent and insignificant results between normal and tumor-exclusive HLAp (data not shown). Since the usage of these three hydrophobic residues is naturally abundant in HLAp, when comparing the immunopeptidome from tissue including normal cells, it is expected that the statistical significance can be weakened and will not be shown as a major difference. Thirteen out of 17 patients carry at least one pΩ W-restraint HLA allotypes (A*24:02, B*4402 or B*44:03) and the usage of W at pΩ was −0.45% in normal-exclusive HLAp and +0.8% in tumor-exclusive HLAp compare to the shared-peptide group (*p* = 0.0185) (Figure 2E, [W]). There was a negative correlation between normal and tumor-exclusive HLAp in the usage of W at pΩ. This indicated the shift of trimming of HLAp in tumor tissue (*r* = −0.788, *p* = 0.0014) (**Figure 2F**). In addition to these two amino acids, the C usage at C-terminus was found to be increased in tumor-exclusive HLAp (+ 0.25%) than that of normal-exclusive HLAp (0.00%) compared to the shared-peptide group (*p* = 0.016) (Figure 2E, [C]). Since there was no clear information available for the binding motif that requires C at pΩ, all 17 samples are included for this calculation. Interestingly, the different C usage of normal and tumor-exclusive HLAp on pΩ may imply the unknown mechanism of HLAp processing under cancer microenvironment.

It has been a long-standing challenge to identify neoantigens directly from small clinical tissues, but our personalized immunopeptidomics analysis overcame that task and successfully revealed the distinct population of immunopeptides in cancer tissue across patients. Intriguingly, despite the long-held theory of “neoantigen depletion” especially with a cancer driver mutation in the late-stage of cancer, the direct identification of neoantigen with a representative cancer driver mutation, such as oncogenic KRAS, from the terminal-stage of colon cancer tissue was achieved by newly established DIM-MS based global-immunopeptidomics approach.

### 2.4. Detectable oncogenic KRAS-carrying neoantigens by global-immunopeptidomics approach

In order to better understand the controversial situation of neoantigen depletion especially from cancer driver mutations, the presentation of neoantigens with oncogenic KRAS mutations was next investigated. Based on the results of global-imunopeptidomics, *i.e.*, for the first time and the direct identification of neoantigen carrying KRAS-G12V from tissue, the cell line Colo-668 was prepared as a positive control carrying the same pair of KRAS-G12V with HLA-A*11:01 as ID 172. Other cell lines, RCM-1, QGP-1 and KP-3 were also prepared as KRAS-G12V carrying cells with different HLA allotypes (**Figure 3A**). First, through global-immunopeptidomics, we analyzed all these cell lines whether any of these cell lines present KRAS-G12V-carrying neoantigens at a detectable level. From 2 of the 4 samples, the neoantigens KRAS-G12V [7-16] and KRAS-G12V [11-19], were respectively identified from Colo-668 (positive control) and RCM-1, together with other unique neoantigens (Figure 3A). Again, neoantigens with oncogenic KRAS were detectable by the established global-immunopeptidomics.

**Figure 3.**
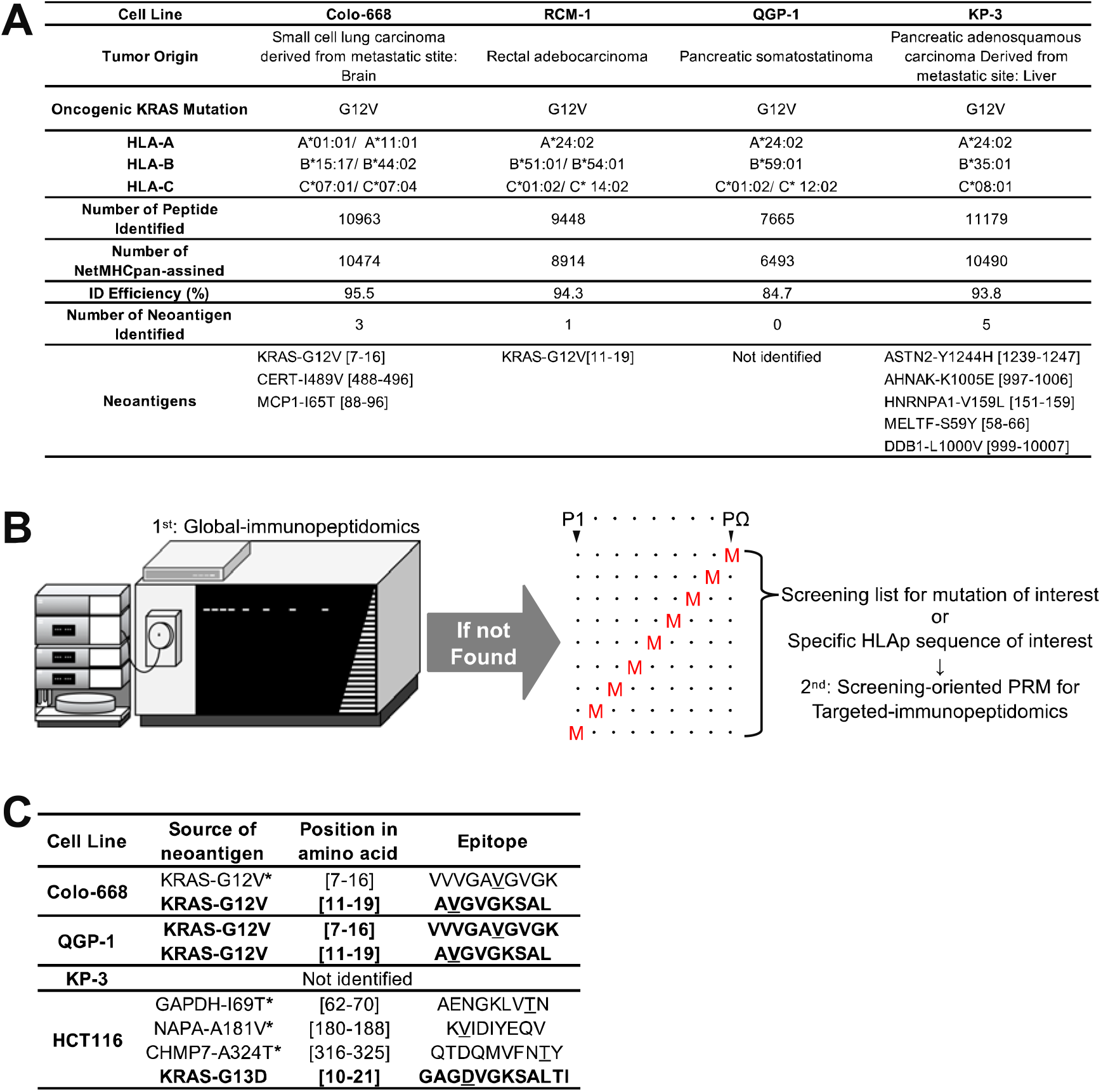
More variations of oncogenic KRAS-carrying neoantigens identified by comprehensive-immunopeptidomics approach. A) Summary of the first screening of KRAS-G12V-carrying neoantigens using cell lines by global-immunopeptidomics alaysis. The cell line Colo-668 was used as a positive control based on the reason for having the genetically identical characteristics of the tumor tissue of ID 172 (carrying HLA-A*11:01 allotype and KRAS-G12V mutation), in which the neoantigen KRAS-G12V [7-16] has been identified. The identification (ID) efficiency was calculated by (number of assigned-peptide by NetMHCpan)/(number of candidate peptides (8 to 15 aa in length)). B) The schematic diagram of comprehensive-immunopeptidomics approach. If the HLAp/neoantigen of interest is undetectable by the in-depth global-immunopeptidomics, then the targeted-immunopeptidomics will be performed by screening-oriented PRM. For the screening of oncogenic KRAS-carrying neoantigens, the inclusion lists containing m/z information of possible sequences of 9 to 12 amino acids in length with KRAS mutation spanning from P1 to PΩ, of charge state 2 and 3, were used. The red M indicates the position of the mutation in the putative HLAp sequence. C) Summary of identified oncogenic KRAS-carrying neoantigens identified by targeted-immunopeptidomics. The asterisk indicates neoantigen that was detectable by firstly performed global-immunopeptidomic analysis of the respective samples. Neoantigens identified were used as positive controls in the PRM. Newly identified oncogenic-KRAS-carrying neoantigens by targeted-immunopeptidomics were written in bold. D) The schematic diagram of comprehensive-immunopeptidomics combining the established global- and targeted-immunopeptidomics approaches. In the global-immunopeptidomics approach, due to the detection limit of the instrument, the ionization efficiency of the HLAp and the ion suppression effect, undetectable potential HLAp/neoantigens will inevitably arise, while that missing information can be complemented, at least in part, by the targeted-immunopeptidomics. Reciprocally, the FAIMS-assisted global-immunopeptidomics approach guarantees the unreachable robust immunopeptidome by targeted-immunopeptidomics.

### 2.5. Targeted-immunopeptidomics analysis revealed more variation of potential neoantigens carrying oncogenic KRAS

The successful detection of neoantigens carrying KRAS-G12V by global-immunopeptidomics drove us to further investigate the potential neoantigens that may have been overlooked in the past. In order to reveal the presentation of potential neoantigens, we prepared an inclusion list for PRM, which contains sequence(s) of HLAp of previously detected neoantigen(s) by global-immunopeptidomics as positive control(s), and combined with the sequences of the HLAp of interest. The predicted m/z of the HLAp sequences of interest, *i.e.*, 9-12 amino acids in length, with KRAS mutation spanning from P1 to PΩ (**Figure 3B**) were included for a screening purpose. By this inclusion list, screening-oriented targeted immunopeptidomics was conducted by PRM. As a result, the positive control, KRAS-G12V [7-16] was again detected by targeted-immunopeptidomics from Colo-668 (**Figure 3C, Figure S6**). And this time, the same neoantigen was also identified from QGP-1 (Figure 3C, Figure S6). In addition, a new sequence of KRAS-G12V [11-19] was detected from Colo-668 and QGP-1 (Figure 3C, Figure S6). Screening-oriented PRM enabled the detection of potential neoantigen of interest and based on these results, established as targeted-immunopeptidomics. Although the knowledge of HCT116 immunopeptidome has been accumulated to date, the neoantigen carrying KRAS-G13D mutation has never been detected by previous global-immunopeptidomics. Therefore, we decided to challenge whether it is possible to identify KRAS-G13D-carrying neoantigens through by established targeted-immunopeptidomics. Here, for the first time, a neoantigen with KRAS-G13D [10-21] was successfully identified from HCT116 (Figure 3C, **Figure S7**). This result indicated that some spectra will inevitably be missed by the efficient global-immunopeptidomics but the missing information can be retrieved, at least in part, through the established targeted-immunopeptidomics. These results indicated that global-immunopeptidomics based on DIM-MS and targeted-immunopeptidomics based on PRM can complement the lack of immunopeptide knowledge, which is unachievable by the unilateral approaches. The fact of previous analyses of immunopeptidome conducted mainly by the global approach with insufficient efficiency, it is suggested more potential neoantigens with cancer driver mutation still remain undetected and that can be identified by the targeted-immunopeptidomics in the future.

## 3. Discussion

In this study, we first established an ideal method to identify the robust immunopeptidome, allowing the comparative analysis of normal- or cancer-exclusive immunopeptidome, that revealed cancer specific signatures of HLAp, such as neoantigens and altered trimming, from a small amount of clinical tissue. Then next, by the established targeted-immunopeptidomics approach, we further demonstrated the presentation of more variation of potential neoantigens carrying oncogenic KRAS.

In the past, immunopeptidomics analysis required a large amount of samples especially from clinical tissues.^[7,9–10]^ Alternatively, patient-derived cancer cells must be prepared as primary cultures or culture organoids to ensure a sufficient amount of sample.^[19–20]^ The purified cultures of patient-derived cancer cells are convenient for various analyses. However, there is a possibility that time-consuming process of cell culture without cancer microenvironment may affect the presentation of HLAp and differ from its original pathological state. Although the mouse xenograft model can also ensure the patient-derived cancer cells, the response to ICIs may vary depending on the host organisms, as shown in the discrepancy result of clinical trial of MEK inhibitor^[21]^ with PD-L1 treatment.^[22]^ Therefore, it is very complicated to evaluate the efficacy of cancer immunotherapy through model materials. The FAIMS-assisted DIM-MS paved the way for the identification of approximately 5,000 of HLAp from 40 mg of tissue (a cube of 3-5 mm in size) without the need for bias-prone chemical prefractionation procedures.^[23]^ Thus, our global-immunopeptidomics approach is an ideal method for direct and robust analysis from scarce samples.

In principle, more mutation load means more possible neoantigens and is considered to be associated with the better outcomes of ICI treatment.^[24–25]^ However, despite the highest mutation load in sample ID 119 (1,752, **Table 2**), no neoantigen was detected even from a total of 6,983 HLAp. According to previous *in vitro* studies, IFN-γ stimulation induces immunoproteasome, which facilitates the peptide trimming in an HLA-Class I-preferred manner, that increases HLAp presentation.^[6, 20]^ Therefore, the presentation of HLAp is considered to be increased in cancer tissue due to the cancer microenvironment under IFN-γ stimulation. However, there have been few reports of direct comparisons of immunopeptidome between normal and cancer from the same individual. In this study, as with mutation load, the number of HLAp identified in cancer tissue was not significantly different from the number identified in normal tissues (Figure S3). These results indicated that because tissue contains various factors that affect HLAp presentation, direct interrogation of tissue samples may not simply recapitulate the previous *in vitro* studies conducted by purified cells.

**Table 2.** Clinical sample information of CRC tissues used in this study.

| ID | Number of Somatic Mutation | Status of Oncogenic KRAS | HLA-A |  | HLA-B |  | HLA-C |  |
| --- | --- | --- | --- | --- | --- | --- | --- | --- |
| No. 51 | 55 | WT | A02:07 | A11:01 | B40:01 | B46:01 | C01:02 | C07:02 |
| No. 60 | 60 | WT | A11:01 | A24:02 | B15:01 | B40:01 | C03:04 | C04:01 |
| No. 78 | 53 | WT | A02:01 | A24:02 | B40:01 | B52:01 | C07:02 | C12:02 |
| No. 84 | 42 | G13D | A11:01 | A24:02 | B54:01 | B59:01 | C01:02 |  |
| No. 103 | 83 | G12V | A24:02 | A26:01 | B40:02 | B52:01 | C03:04 | C12:02 |
| No. 107 | 74 | G12D | A02:01 | A24:02 | B15:01 | B40:02 | C03:04 | C08:03 |
| No. 112 | 110 | G12D | A02:01 | A26:01 | B15:01 | B55:02 | C01:02 | C04:01 |
| No. 119 | 1752 | G13D | A24:02 | A33:03 | B44:03 | B55:02 | C01:02 | C14:03 |
| No. 172 | 123 | G12V | A11:01 | A24:02 | B35:01 | B40:02 | C01:02 | C03:03 |
| No. 229 | 31 | WT | A24:02 | A33:03 | B15:18 | B44:03 | C07:04 | C14:03 |
| No. 241 | 32 | WT | A24:02 | A33:03 | B40:01 | B40:06 | C03:04 | C12:02 |
| No. 256 | 61 | G12D | A02:01 | A02:06 | B35:01 | B38:02 | C03:03 | C07:02 |
| No. 257 | 61 | G12V | A02:01 | A24:02 | B15:01 | B52:01 | C03:03 | C12:02 |
| No. 258 | 34 | G12D | A24:02 | A33:03 | B44:03 | B52:01 | C12:02 | C14:03 |
| No. 259* | 56 | WT | A02:01 |  | B44:02 | B55:02 | C01:02 | C05:01 |
| No. 260 | 14 | WT | A02:07 | A24:02 | B46:01 | B54:01 | C01:02 |  |
| No. 261* | 69 | G12A | A11:01 | A24:02 | A24:02 | B13:01 | B52:01 | C07:02 |

\* Liver metastasized tissue

In previous immunopeptidome analyses using *in vitro* samples with or without IFN-γ stimulation, it has been reported that the chymotryptic trimming (mainly by L, I, V) becomes dominant in response to the induction of immunoproteasome.^[7,19]^ However, of the amino acids targeted by chymotrypsin, only the tryptophan (W) trimming was statistically increased in the cancer-exclusive HLAp in this study. Intriguingly, it has been reported that chronic exposure to IFN-γ induces the skipping/frameshifting translation at certain tryptophan codons that results in translation of aberrant peptides. In addition, these aberrant peptides have been reportedly to become a source for neoantigens in melanoma cells.^[26]^ The association between the increase of W-trimming in cancer-exclusive HLAp in this study and the translation of aberrant peptide is of interest, and this shall be clarified by the customized database search^[26]^ or the *de novo* MS analysis in future.^[8,27]^

Recently, the control immunopeptidome, *i.e.*, HLAp presented in normal cells, has been recognized as indispensable for more refined analysis in cancer immunopeptidomics.^[28]^ In addition to the curated immunopeptidome established from multiple genetic backgrounds, the dataset of personalized immunopeptidome established from identical individual is also worthwhile to elucidate the new insights by keeping the complex factors that influence the HLA presentation as simple as possible. Although further analysis of various pathological conditions is required to delineate the new aspects of processing/presentation of HLAp, the robust dataset of individual immunopeptidome from both normal and cancer obtained in this study will be of use for those who want to explore the new landscapes in immunopeptidomics.

Due to the eradication of neoantigen-presenting cancer cells by anti-tumor immunity,^[11–12]^ it has been suggested that the frequency of neoantigen presentation on solid tumors might be low,^[5,10,19]^ especially in CRC.^[2,29]^ However, in this study, 2 neoantigens, one of which carried KRAS-G12V, were directly identified from stage IV CRC tissue. sRecently, the affinity of two KRAS-G12V-carrying neoantigens for HLA-A*03:01 or A*11:01 have been quantitatively characterized by PRM using KRAS-G12V-transduced monoallelic cell samples.^[30]^ Although the monoallelic expression of HLA is a useful experimental model to characterize the affinity and the restraint of HLAp, the actual presentation of HLAp may vary due to the similarity of binding motifs between intrinsic HLAs. In this study, the established targeted-immunopeptidomics enabled the direct detection of neoantigen of interest under full-allelic conditions. The newly identified neoantigens that carry oncogenic KRAS (G12V or G13D) were not predicted as possible-binders by NeMHCpan4.1. Still, judging by the motif viewer of NetMHCpan, due to the tolerable amino acids for the anchor positions, the possibility of KRAS-G12V [11-19] binding to the HLA-B*15:17 in Colo-668 as well as to the HLA-C*12:02 in QGP-1 seemed undeniable. As for the possibility of the presentation of KRAS-G12V [7-16] in QGP-1, it has previously been clearly shown that valine at p2 and lysine at pΩ are preferred in HLA-C*12:02-bound HLAp.^[31]^ The feature of this binding preference is fully in line with the amino acid sequence of KRAS-G12V [7-16], still this peptide sequence has not been predicted as a possible binder. These results indicated that the incomplete prediction algorithm will eventually erroneously predict some possible-binders as non-binders. Owing to the wide variety of binding motifs of a vast number of HLA types, there is no wonder that it will inevitably require more time to build a complete prediction algorithm. In the analysis of HCT116, the KRAS-G13D-carrying neoantigen was detectable only by the targeted-immunopeptidomics but not by the repeated robust global-immunopeptidomics. This implies that it is very likely that most of the potential neoantigens carrying cancer driver mutations have yet to be identified. Even without information of actually-presented neoantigens, the immunotherapy by autologous T cells that recognize KRAS-G12D-carrying neoantigen in lung-metastasized CRC has been reported as effective and successful in the previous clinical trial.^[32]^ However, it takes extra time and cost to screen the immunoreactive T cell clones, and that will become an obstacle to establish more versatile cancer immunotherapy. In addition to discovering more unknown potential neoantigens, the direct confirmation of the presentation of HLAp/neoantigen of interest indeed in the aimed subject should further contribute to the more effective decision-making. Thus, the established global- and targeted-immunopeptidomics approaches shown here grant the best knowledge by making the most of precious clinical samples and that will enlighten us new outlook for the construction of sophisticated prediction algorithms, recurrent neoantigen search and the suitable epitope selection for immunotherapy in precision cancer medicine. This advances in comprehensive-immunopeptidomics approach holds promise to open a next phase of cancer immunology with new prospects from both basic and clinical aspects.

## 4. Experimental Section

### Cell line and clinical tissue samples

The cell lines used in this study are described in **Supporting Information.** The clinical information including sample ID, mutation load, oncogenic KRAS status, and the genetic background of class-I HLA are listed in Table 2. All surgically removed clinical tissues was dissected into the desired size and immediately cryopreserved at least once before use. Approximately 40 mg of net weight tissue was subjected to immunoaffinity purification.

### Ethics approval

Metastatic colon cancer tissues at pathological stage IV and its adjacent normal tissues were obtained from patients who all had provided written informed consent for this study. This study was approved by the ethical committee of the Japanese Foundation for Cancer Research (JFCR) (Ethical committee number 2010-1058).

### Immunoprecipitation of HLA-Class I complex and HLAp purification

The in-house purified anti-panHLA antibody (W6/32) was used for immunoprecipitation of HLA complex. Detailed information of sample preparation for immuopeptidomics can be found in Supporting Information. Briefly, desired amount of cell pellet or tissue were lysed by 1 ml of lysis buffer containing [20 mM] HEPES, [150 mM] NaCl, 1% NP-40, 0.1% SDS and 10% Glycerol on ice. After centrifugation for clarification, the supernatant was subjected to immunoprecipitation for overnight at 4 °C on a rotating rotor. After the washing steps, the trimer of HLA complex was dissociated and eluted by 500 μl of 1% TFA. The obtained eluate was further processed by tC18 SepPak (Waters), and the HLAp was enriched by 500 μl of 20% acetonitrile. The prepared HLAp samples were then dried by an evaporator and stored at −30 °C until MS analysis.

### Construction of customized database for cell lines

The customized databases for cell line was established by whole exome sequencing (WES) analysis from the COSMIC Cancer Cell Line Project. All listed somatic mutations were processed by in-house built pipeline as previously reported.^[33–34]^ Then, the obtained cell line-specific mutations were combined respectively with Swiss-Prot human proteome database (20,181 entries) from UniProt website to perform the customized database search.

### HLA typing, WES and in-house pipeline for personalized databases of clinical tissues

For clinical tissue samples, genomic DNA was extracted by QIAmp DNA mini Kit (QIAGEN) from normal and tumor tissues. To establish the personalized database for global-immunopeptodomics, WES analysis was performed in all samples as previously described.^[34–35]^ The cancer-specific mutations were extracted by subtraction of non-relevant mutations identified in normal tissue. For more details on the construction of the personalized database, please refer to the Supporting Information.

### Global-immunopeptidomics by high-field asymmetric-waveform ion mobility spectrometry (FAIMS)-assisted differential ion mobility (DIM) mass spectrometry (MS)

The sample includes HLAp was first trapped by precolumn (C18 Accleim PepMap 100 C18 Trap Cartridge, Thermo Fisher Scientific) and then separated by analytical column (Aurora UHPLC Column (C18, 0.075 × 250 mm, 1.6 um FSC, ESI, ion opticks) coupled with a nanospray Flex ion source for electrospray ionization (Thermo Fisher Scientific), were used through an Ultimate 3000 RSLC nano HPLC system at a flow rate of 200 nl/ min using a linear gradient starting from 2% solvent B (0.1% Formic acid in acetonitrile) to 28 over 55 min, a 1 min hold at 95% Solvent B a prior to a 1 min analytical column equilibration with 2% B. In order to validate the benefit of FAIMS applied DIM-MS in immunopeptidomics, liquid chromatography tandem mass spectrometry (LC-MS/MS) and LC-FAIMS-MS/MS conditions were firstly compared. For LC-FAIMS-MS/MS, the FAIMS-Pro interface (Thermo Fisher Scientific) was installed onto an Orbitrap Fusion Lumos Tribrid mass spectrometer and operated with default parameters except for the compensation voltage (CV) settings for gas-phase fractionation.^[36]^ For LC-MS/MS, we uninstalled FAIMS device from mass spectrometer and performed analyses. Under LC-FAIMS-MS/MS condition, sample was seamlessly fractioned by 3 CVs per single analysis. Total 3 CV sets were used per sample and each CV set includes as follows; set1 (CV = −40, −60, −80 V), set 2 (CV = −35, −50, −70 V) and set 3 (CV = −45, −55, −65 V). Therefore, total 3 analyses (raw data), were acquired per sample. The parameters for mass spectrometry were optimized for immunopeptidomics with small amount of sample. Full MS (rage from 320 to 850 m/z) in the Orbitrap were acquired at resolution of 60 k followed by a MS2 acquisition at resolution of 15k. Maximum injection time for full MS scan was 50 ms with an auto gain control (AGC) of 4e5. Maximum injection time for MS2 scan was 100 ms with an AGC 1e4 followed by top speed MS2 acquisition by the ion trap. Charge state 2 and 3 were selected for fragmentation by collision-induced dissociation (CID) at rapid scan rate at 30% collision energy. The easy-IC system was used for the internal calibration lock mass during the data acquisition. The HCT116 samples were prepared from 1e8 cells and proportionally injected into MS to match the indicated number of cells. For the analyses of 1e8 of HCT116 cells and clinical tissue samples, firstly 1/ 20 amount of sample volume was used for analyses of 3 CV sets respectively to check sample conditions, and then from the remaining, 5/ 20 amount of sample volume was used for 3 CV sets respectively. Therefore, total 6 analyses were performed per clinical tissue sample. All acquired files were searched against the corresponding customized/personalized database. The LC/MS raw data, summarized result files and the list of identified HLAp from samples can be found at a public proteomic database, Japan Proteome Standard Repository/Database (jPOST) as follows; HCT116 HLAp without FAIMS in JPST001072, HCT116 HLAp with FAIMS in JPST001066, Global identification of HCT116 HLAp in JPST001068, HLAp from normal regions of CRC tissues in JPST001070, and HLAp from tumor regions of CRC tissues in JPST001069.

### Database search for global-immunopeptidome

The obtained raw data was processed by Proteome Discoverer version 2.4 using Sequest HT engine against corresponding database. For the enzymatic designation, the search was set to no-enzyme (unspecific). The precursor mass tolerance was set to 10 ppm, and the fragment mass tolerance was set to 0.6 Da. The maximum number of missed cleavages was set to zero. Methionine oxidation and cysteine carbamidomethylation were included as dynamic modifications. For the percolator filtering, the false discovery rate (FDR) was set to 0.01 (Strict) to identify highly confident peptides. The output “peptide groups” was exported as a list in Excel format. The peptide sequence in the list were first deduplicated only for unique sequences, and then filtered by peptide length from 8- to 15-mer as a candidate-HLAp. Finally, the sequences predicted as non-binding by NetMHCpan 4.0^[37]^ were further excluded from the list to determine the assigned-HLAp. Simultaneously, for unsupervised clustering of HLAp and motif sequences, Gibbs Cluster 2.0^[38]^ and Seq2 logo^[39]^ were used to confirm whether the identified HLAp/motifs was consistent with the HLA allotypes according to each sample.

### Profiling of HLAp in normal and tumor tissues

In order to investigate the possible distinctions between normal and tumor-derived HLAp, the intersection of HLAp between normal and tumor tissue was calculated and drew Venn diagrams by publicly available website (http://bioinformatics.psb.ugent.be/webtools/Venn/). Based on the Venn diagrams, the personal immunopeptidome was classified into the following 3 groups: (1) normal-exclusive, (2) shared in both normal and tumor tissues (here called “shared-peptide”) and (3) tumor-exclusive. The frequency of the usage of each amino acid at the C-terminal of HLAp (pΩ) was calculated as a percentage (%) relative to the total number of HLAp in each group. The difference in the frequency of usage of the amino acid at the C-terminus between the normal-exclusive group and the tumor-exclusive group was calculated by comparing with the frequency of the shared-peptide group, respectively.

### Targeted-immunopeptidomics by screening-oriented parallel reaction monitoring (PRM)

The oncogenic KRAS status for cell lines were available by “THE RAS INITIATIVE” at National Cancer Institute (NCI) of National Institute of Health (NIH) (https://www.cancer.gov/research/key-initiatives/ras/outreach/reference-reagents/cell-lines). The preparation of HLAp samples and the condition of instruments were the same as the above-mentioned global-immunopeptidomics, except for the condition of uninstalled FAIMS-Pro interface. For the data acquisition of the targeted-immunopeptidomics, the PRM strategy was introduced with minor modifications.^[40]^ Briefly, collision CID and the detector ion trap were used at the rapid scan mode. The resolution was set to 15,000 and the maximum ion inject time for 300 ms. To prepare the inclusion list, the predicted m/z was calculated by ChemCalc (https://www.chemcalc.org/peptides)^[41]^ for the HLAp of interest. Obtained raw data was searched against the database that includes the full-length amino acid sequence of corresponding oncogenic KRAS and the source protein of the positive controls by proteome discoverer 2.4. The filtering was performed by target-decoy search^[42]^ with 1% FDR.

### Statistical Analysis

All data were expressed as mean ± standard error (SE). Student’s t-test was used for statistical analysis. The Pearson correlation-coefficient was used to assess the correlation. Statistical significance thresholds were determined at *p* < 0.05 (*), *p* < 0.01 (**), *p* < 0.001 (***) values. Pearson correlation-coefficient was used to assess the correlation.

## Supporting Information

Supporting Information is available from the Wiley Online Library or from the author.

## Acknowledgements

This work was supported by “the Development of Technology for Patient Stratification Biomarker Discovery” of the Japan Agency for Medical Research and Development (20ae0101074s0302) and “Grant-in-Aid for Scientific Research (C)” of Japan Society for the Promotion of Science (20K05759).

---

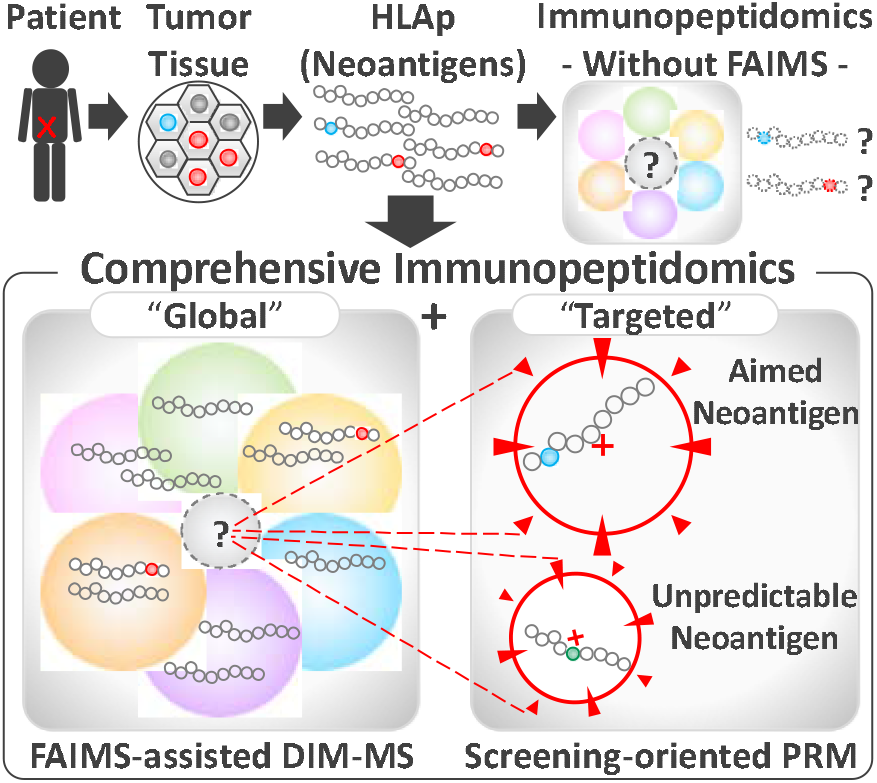
Table of Contents.

## Table of Contents

By this comprehensive-immunopeptideomics analysis, in addition to vigorously expanding the scope of identification of immunopeptides from scarce clinical samples, a screening strategy of neoantigen by mass spectrometry was introduced. Successful direct identification of neoantigen from cancer tissue by mass spectrometry endorse the actual presentation of those neoantigens, which are an optimal target for cancer immunotherapy. This advanced immunopeptidomics analyses will contribute to make cancer immunotherapy more versatile and accessible.

## Supporting Methods

### Cell line samples

The cell lines used in this study are: HCT116 (ATCC), Colo-668 (Sigma Aldrich), RCM-1 (JCRB), QGP-1 (JCRB) and KP03 (JCRB). All cells were cultured in RPMI1640 medium (WAKO) with 10% heat-inactivated fetal bovine serum (FBS), penicillin G and streptomycin, and maintained under the general condition. Cells were washed once with PBS before harvest. Desired number of cell pellets were stored at −80 °C until use.

### In-house purification of antibody and the IP-beads preparation

A hybridoma clone (HB95) of anti-panHLA alpha chain antibody (W6/32) was obtained from ATCC. Hybridoma cells were first expanded in 10% FBS containing DMEM and then cultured in CELL Line flask (WHEATON) with COS medium (COSMO BIO) for antibody production. After 1-week of cultivation, the culture media was collected and affinity purified by Protein A Fast flow (GE). Small amount of eluates were electrophoresed and then stained by GelCode Blue Stain Reagent (Thermo Fisher Scientific) to check IgG fractions. The IgG enriched fractions were pooled and then dialyzed to replace the buffer into PBS. The antibody concentration was examined by BCA assay. To confirm the sufficient affinity of our in-house purified W6/32 antibody against HLA molecules, we tested all batches of W6/32 by general IP and Western blotting by antibody anti-pan HLA class I α-chain molecules (clone EMR8-5, MBL) and β-2-microglobulin (β2M) antibody (PROTEINTECH) in advance. High affinity confirmed W6/32 antibody (800 ug) was then cross linked onto 200 ul of slurry of Protein G Sepharose (GE) by DMP (20 mM) in HEPES (pH 8.4) for 1 hour. After quenching by TBS-T for 1 hour, beads were used for IP or stored at 4 °C before use.

### Immunoprecipitation of HLA-Class I Complex and HLAp purification for Immunopeptidomics

Desired cell pellet was lysed by 1 ml of lysis buffer containing [20 mM] HEPES, [150 mM] NaCl, 1% NP-40, 0.1% SDS and 10% Glycerol on ice. For tissue samples, disposable homogenizer Bio Masher System (nippi) was used firstly with 500 μl of lysis buffer for approximately 40 mg of tissue on ice and then add another 500 μl of lysis buffer to make sample volume as 1 ml. Both cells and tissues samples were cleared by centrifugation at 4 °C, 15,000 rpm for 15 min, the supernatant was collected and conducted immunoprecipitation for 4 °C overnight on rotating rotor. For thorough washing, Bio Spin Column (BioRad) was used afterwards. The washing steps were as follows: 1 ml of lysis buffer for 5 times, 1 ml of PBS-T for 5 times, 1 ml of PBS for 5 times, 1 ml of high-salt PBS for twice, 1 ml of PBS for 5 times and then 1 ml of mass spectrometry-grade water for twice.

The trimer of HLA complex was dissociated and eluted by 500 μl of 1% TFA from IP-beads. Obtained HLA enriched eluate was further processed by tC18 SepPak. For class I HLA peptide, 500 μl of 20% acetonitrile (ACN) was firstly used for elution. HLA peptide including eluates were then dried up by evaporator. To elute the other components of HLA complex to confirm the IP-efficiency, 500 μl of 80% ACN was used for second elution and the obtained eluates were dried up by evaporator as well. Sample purity was assessed by silver staining by Silver Quest (Thermo Fischer Scientific).

### HLA typing, whole exome sequencing and pipe lines for tailored protein database

For clinical tissue samples, genomic DNA was extracted by QIAmp DNA mini Kit (QIAGEN) from normal and tumor tissues. To establish the personalized database for neoantigen identification by immunopeptodomics, whole exome sequencing (WES) was conducted to all samples. The genome libraries for WES were established from the DNA samples using xGen Exome Research Panel (IDT, Coralville, IA), according to the manufacturer’s instructions. The exons were sequenced as 150 base pairs paired-end reads by a NovaSeq6000 system (Illumina, San Diego, CA). The sequence data obtained were then analyzed to select possible germ line variations and somatic mutations. In short, sequences of the whole exome were compared with a reference human genome (hs37d5), and the possible germ line variants and the somatic mutations were identified. The variants in public databases (> 1%) were also excluded. Then cancer somatic mutations were extracted by subtraction of germ line variation identified in normal tissue from tumor tissue respectively. Among those mutations, only the missense mutations were selected and translated into full length of protein and added to the same Swissprot/ Uniprot human database described above as personalized database. The established personalized data base was used for both normal and tumor immunopeptidomics corresponding patient’s samples. The OptiType was employed for 4-digit HLA typing from deep sequencing data. The genetic details including mutation burden, oncogenic KRAS mutations and the HLA genotypes of clinical samples can be found in Table 2.

---

**Figure S1.**
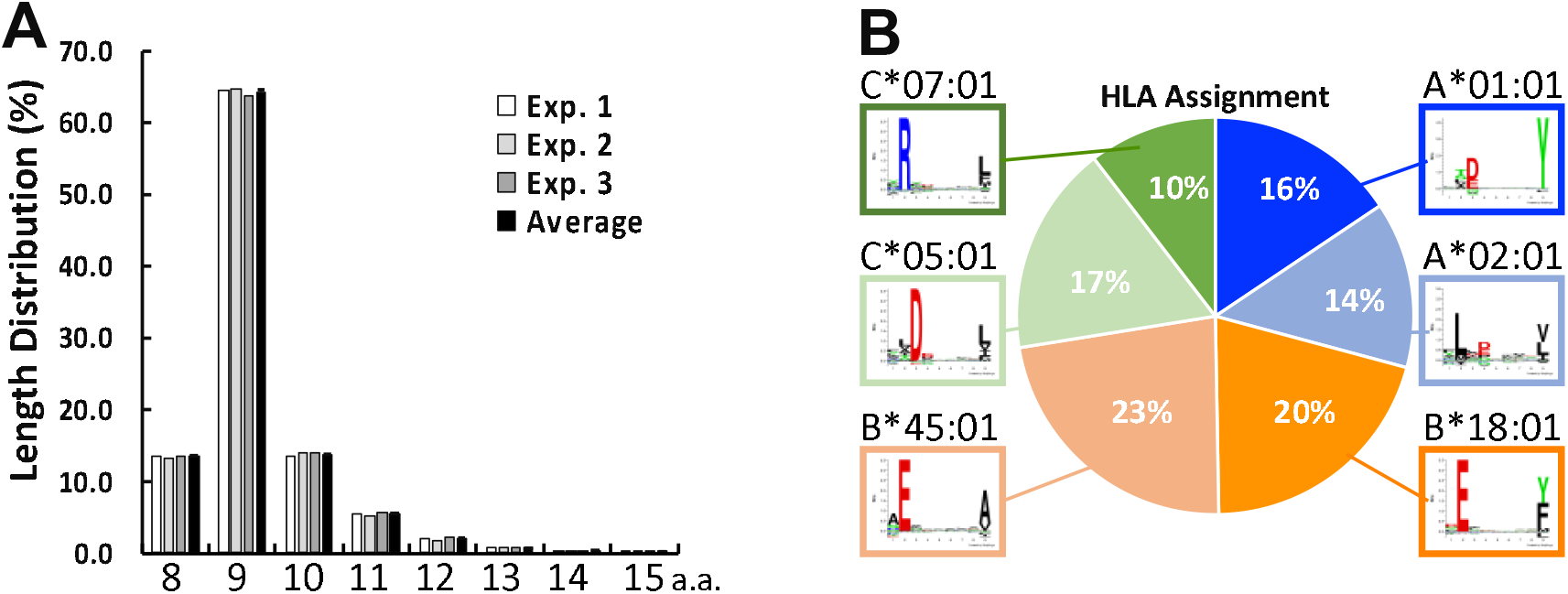
Characteristics of immunopeptidome of a colon cancer cell line, HCT116. A) The length distribution of immunopeptides identified from HCT116. The length of immunopeptides identified in figure 1D was counted. B) Representative image of unsupervised clustering of HLA motif and the population of HLA assignment in the context of HLA allotypes.

**Figure S2.**
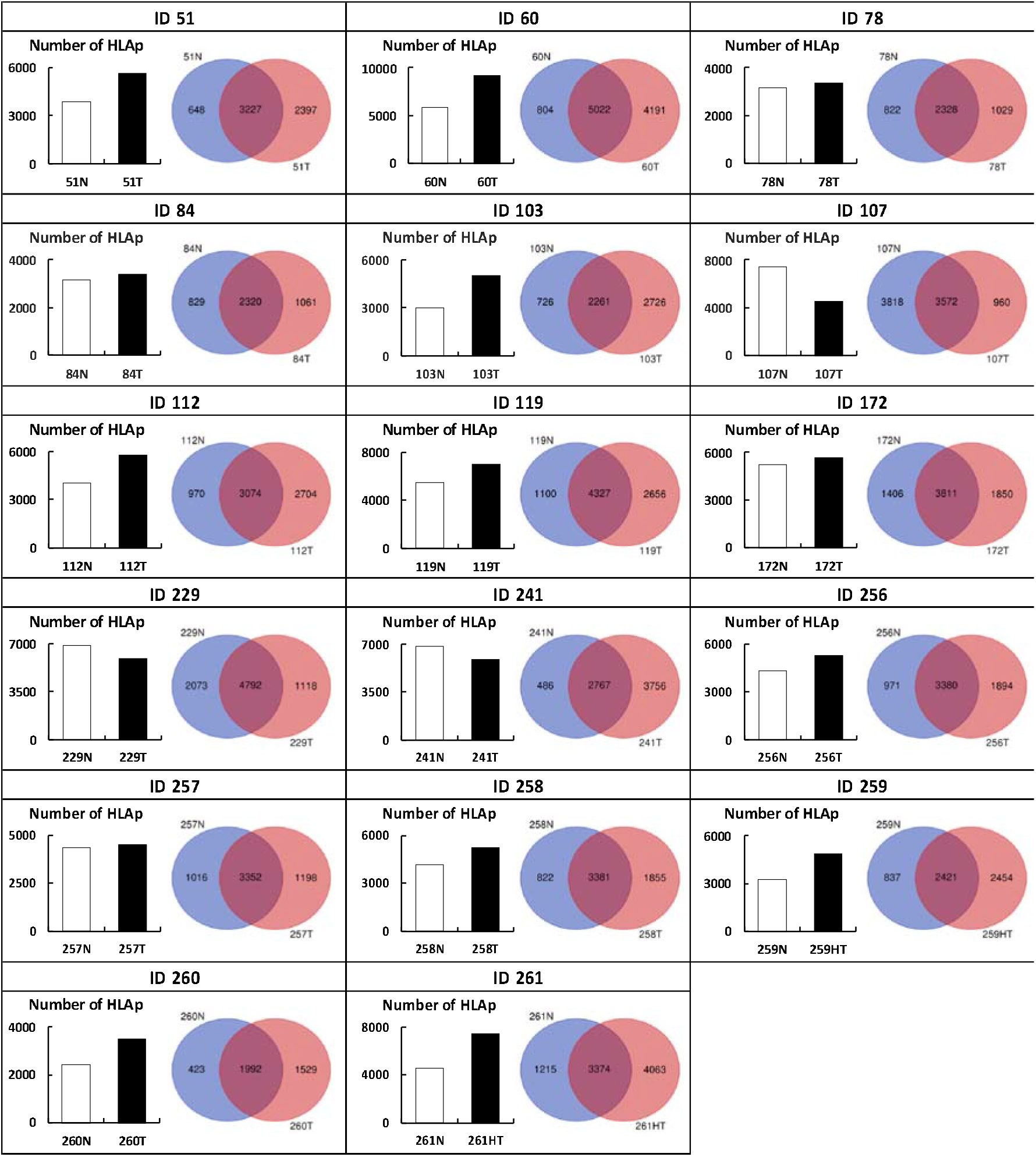
Number of Immunopeptides per sample, per patient. The panel shows the details of identified number of immunopeptide in each normal and tumor tissue samples and the overlap of immunopeptides between normal and tumor was calculated by Venn diagram per person. Venn diagram does not reflect the sample size in this figure.

**Figure S3.**
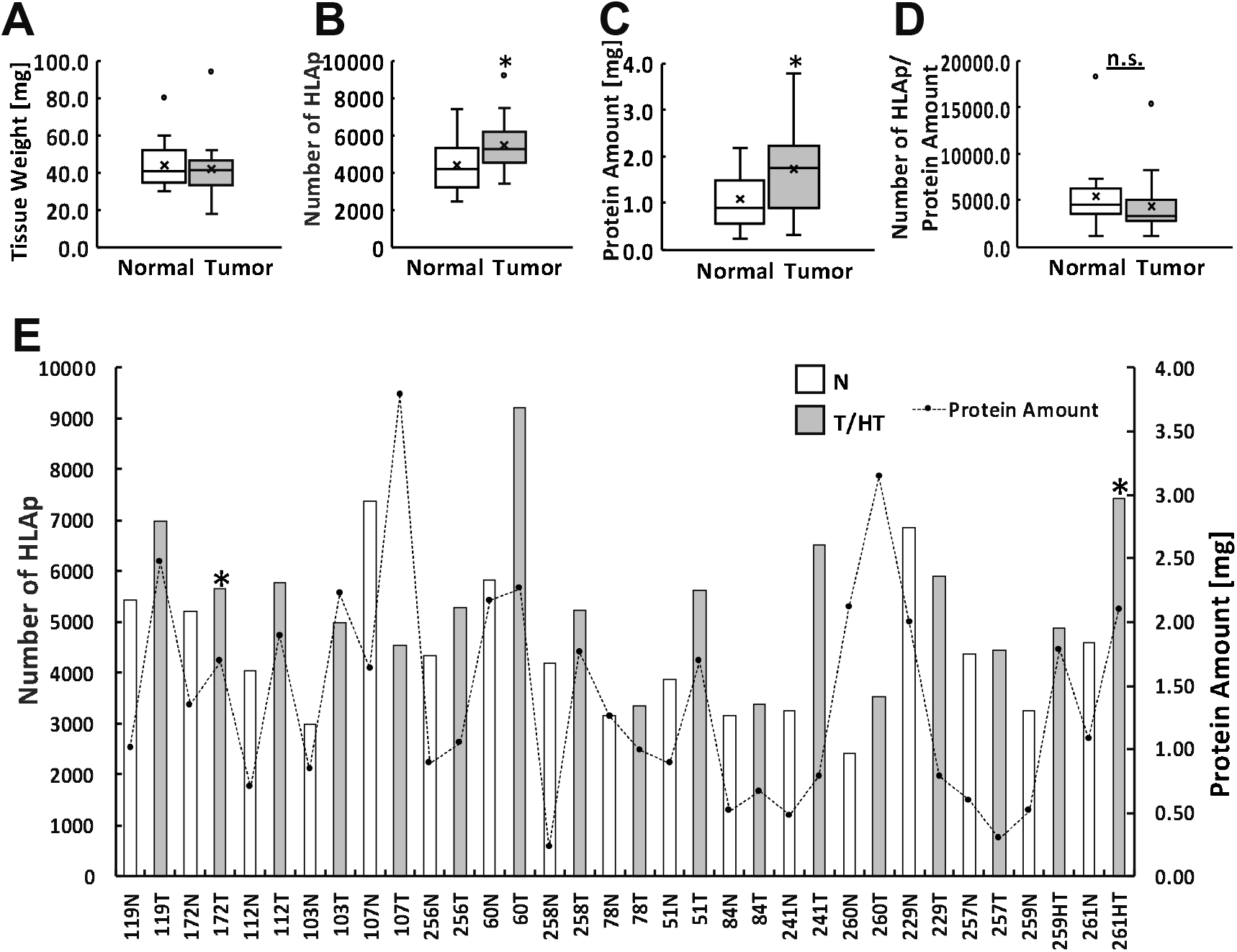
Associations of fundamental sample conditions that may affect the HLAp identification, and the comparison of pathologically-independent immunopeptidome. A) A box plot depict a net weight of tissues, B) a total number of identified immunopeptides, C) a protein amount included in lysate, and D) the number of identified immunopeptides normalized by protein amount. The apparent significance of increased HLAp in tumor tissue than normal tissue was cancelled when normalized by protein amount in the lysates. E) The composite graph of depicts the number of immunopeptide identified in each sample (white bars indicate normal tissues, gray bars indicate tumor tissues) and the amount of protein used in this study. The asterisk indicates the sample in which neoantigen has been identified. There was no consistent association between the number of HLAp identified and the amount of protein used, or the neoantigen detection.

**Supplementary Figure 4.**
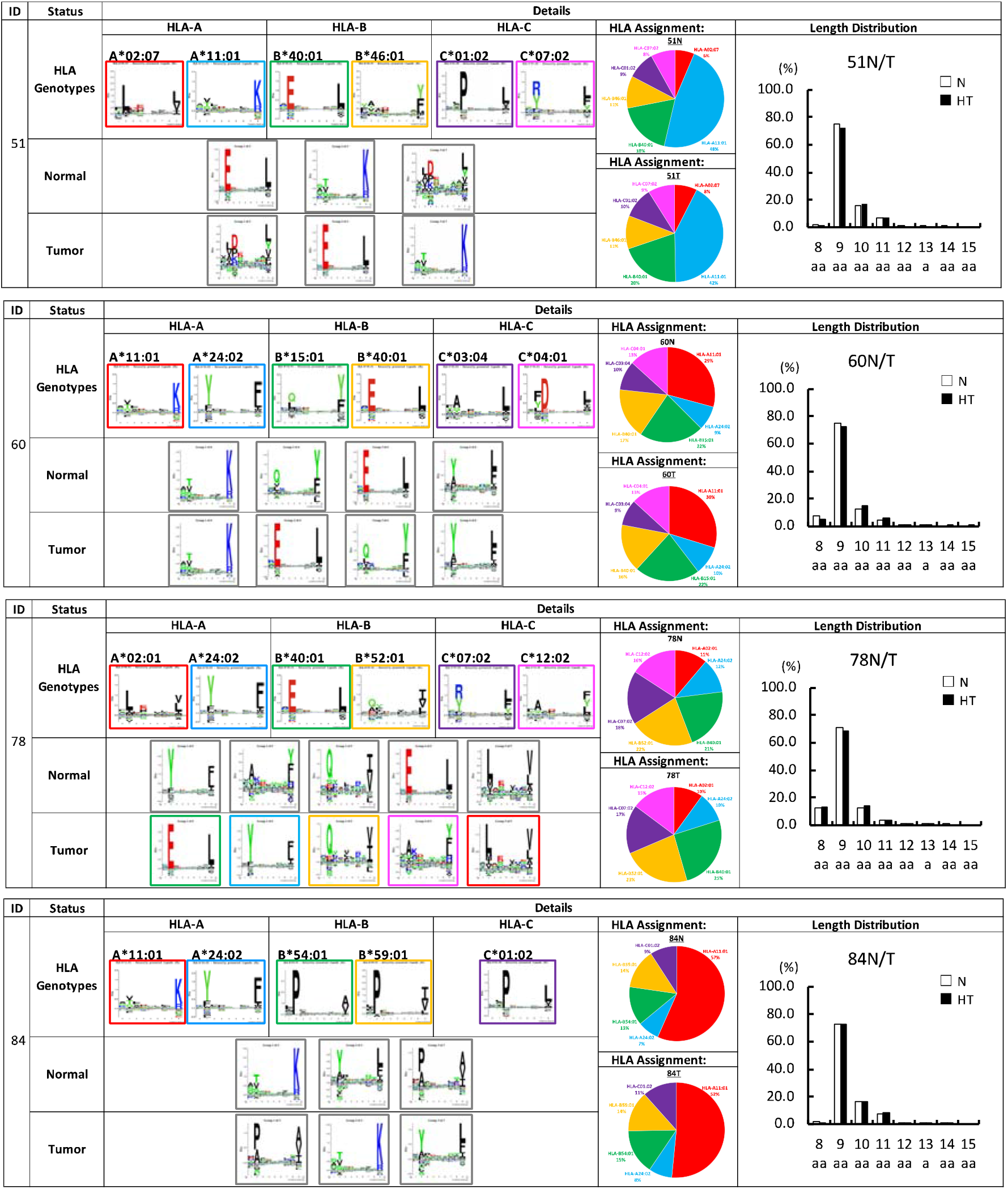
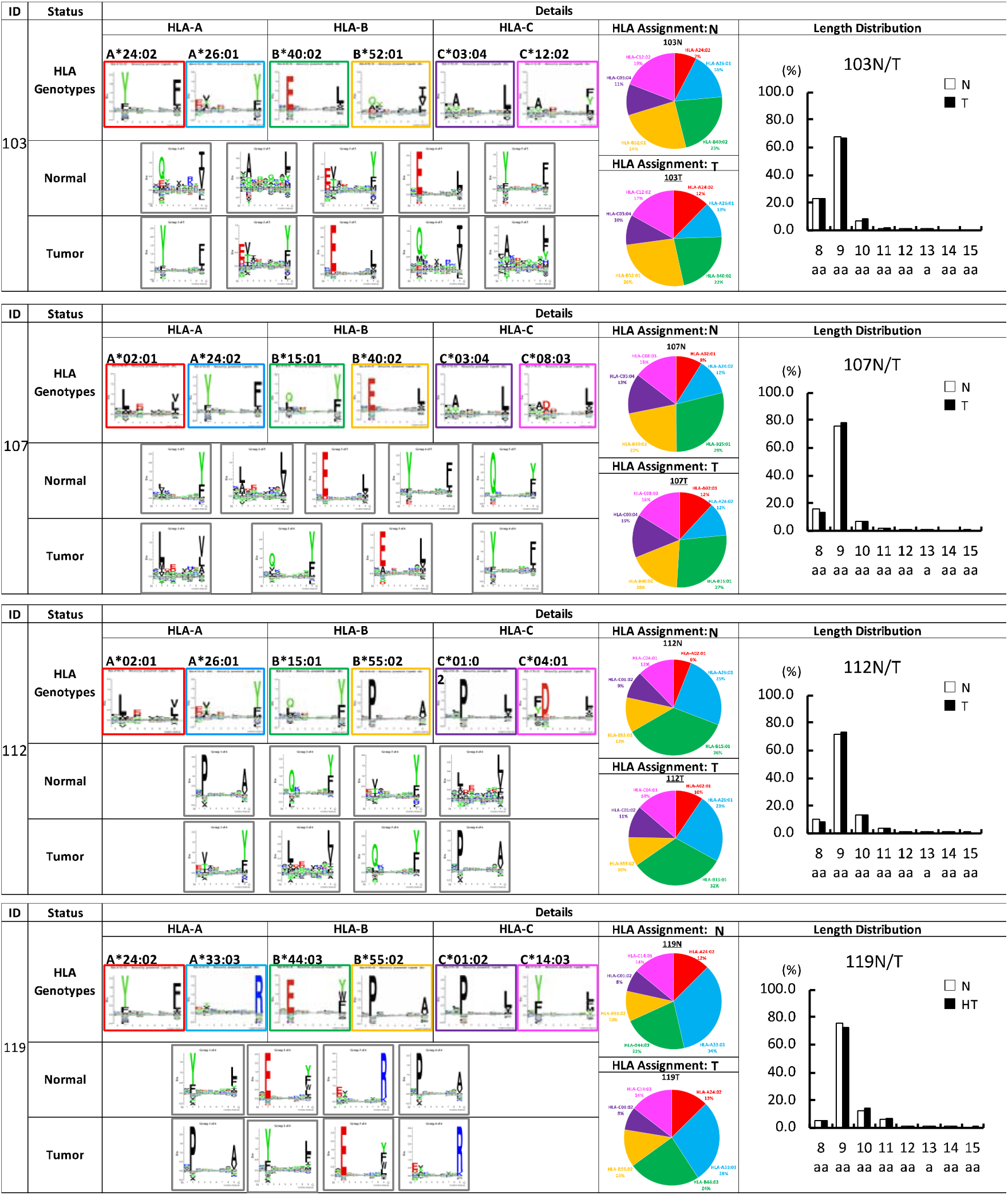
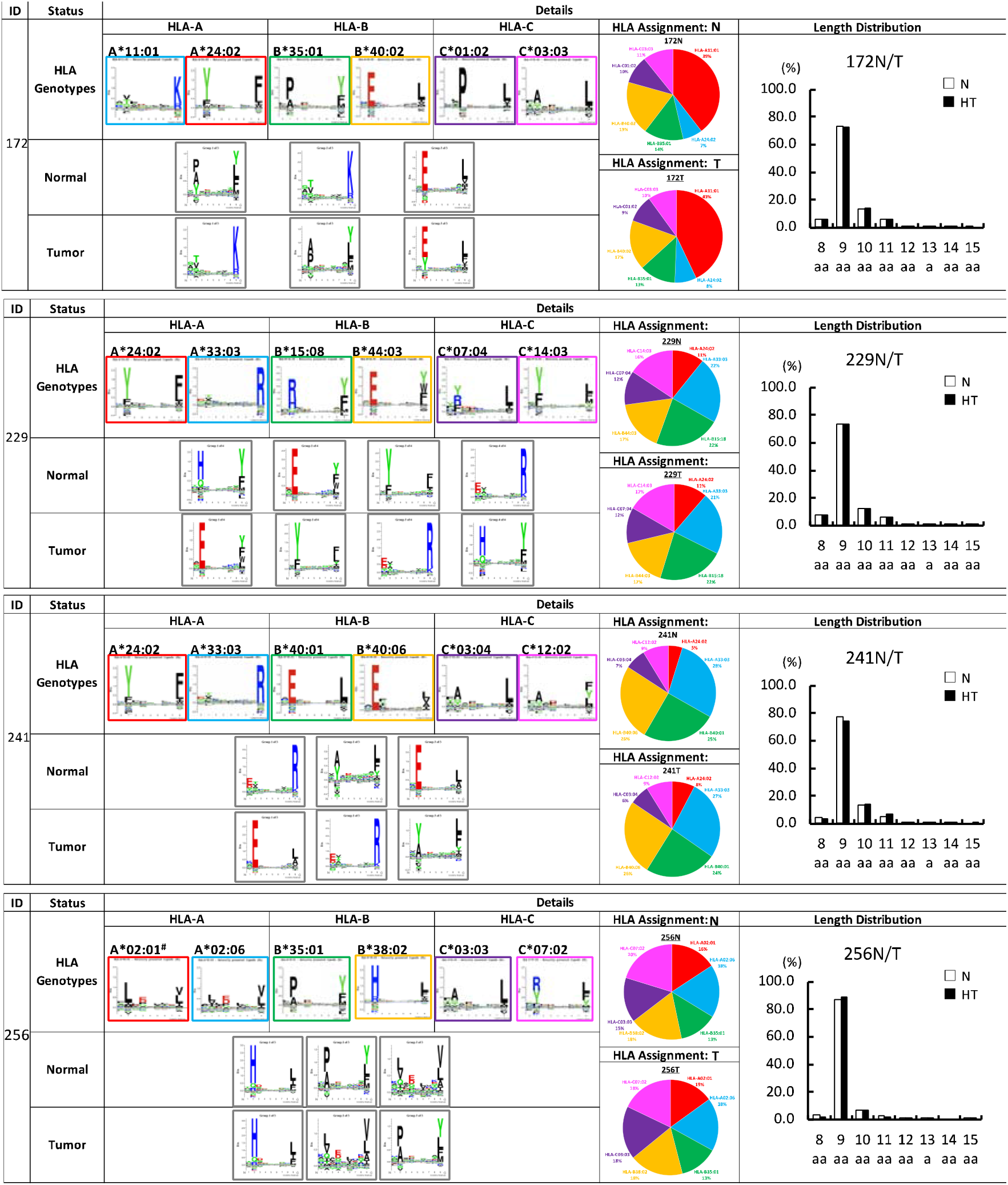
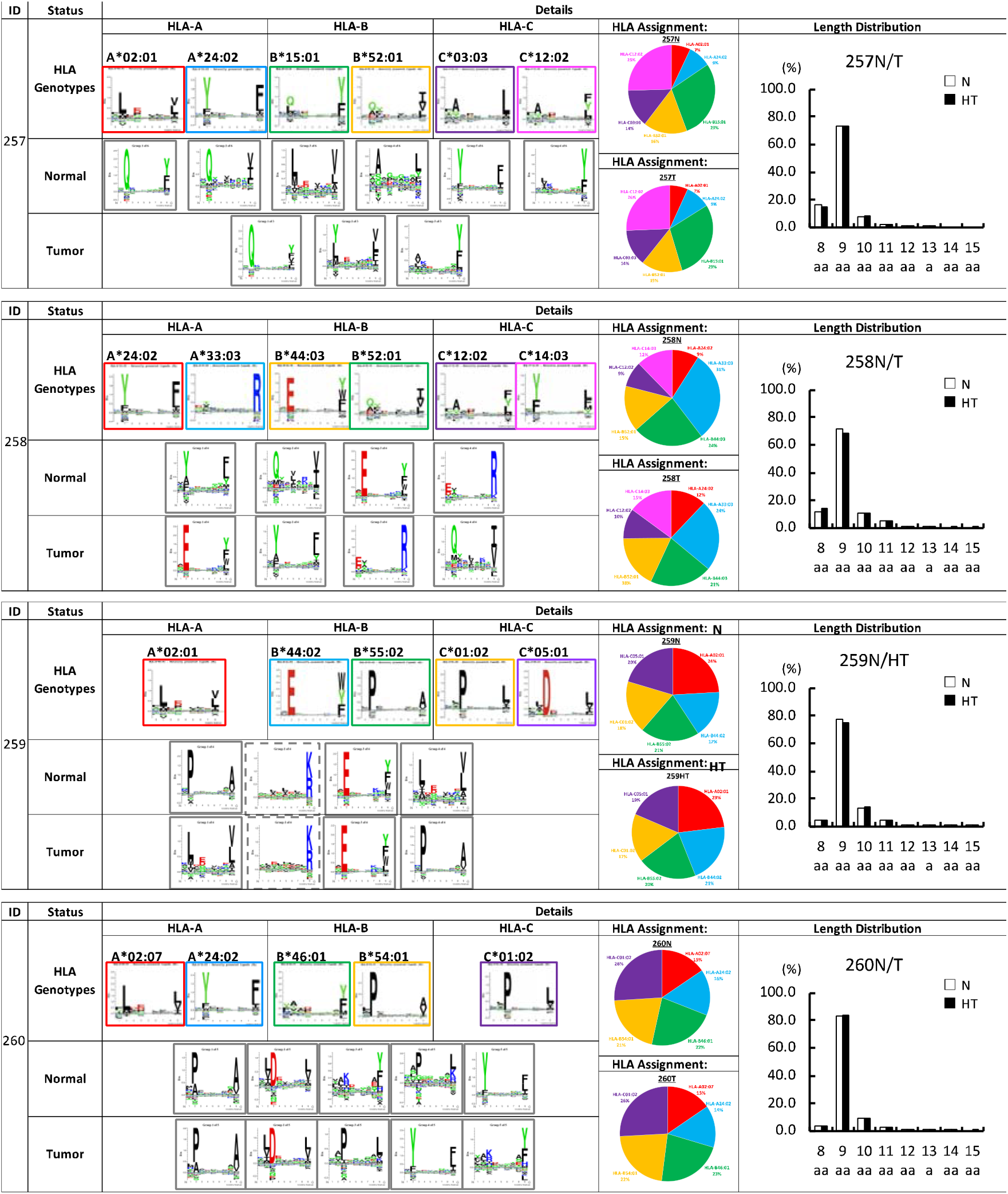
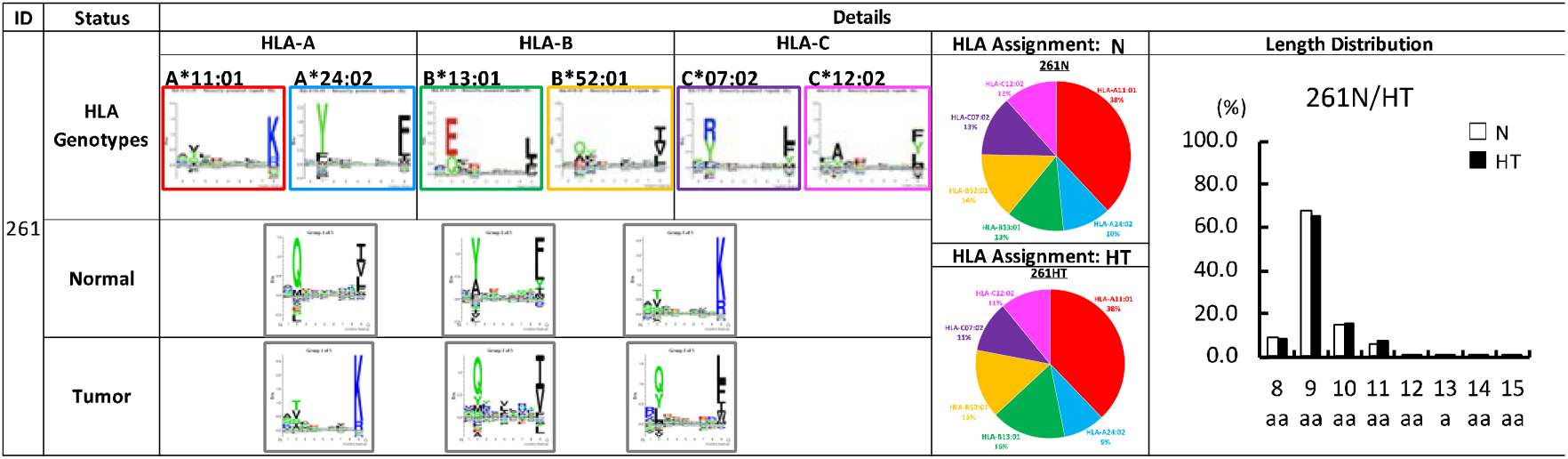
Comparison of unsupervised clustering, assignment and the length of HLAp in individual immunopeptidome. HLA allotype and its motif sequence, unsupervised clustering of actually identified immunopeptides, immunopeptide assignment to corresponding HLA allotypes, and the length distribution of immunopeptides are shown individually. The length of immunopeptides were compared between normal and tumor tissues in identical patients. In these parameters analyzed, there were no differences between normal and tumor immunopeptidome in each patients.

**Figure S5.**
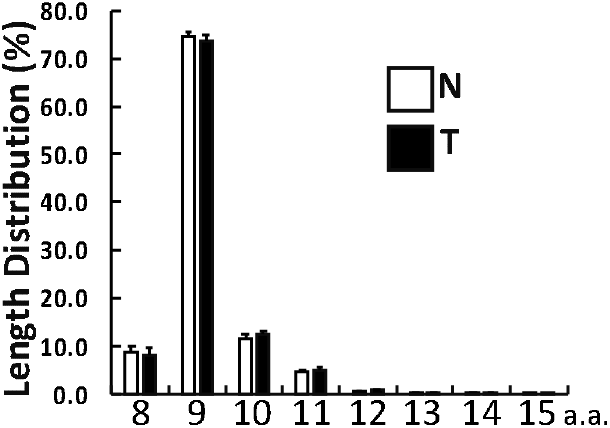
Comparison of personalized immunopeptidome analysis for CRC tissue samples. The length distribution of immunopeptides (44,785 HLAp in total, as shown in Figure 2A). Nine amino acids in length dominated more than 75% of overall immunopeptidome. Eight to 12 amino acids in length covered more than 99.7% of overall immunopeptidome in this study.

**Figure S6.**
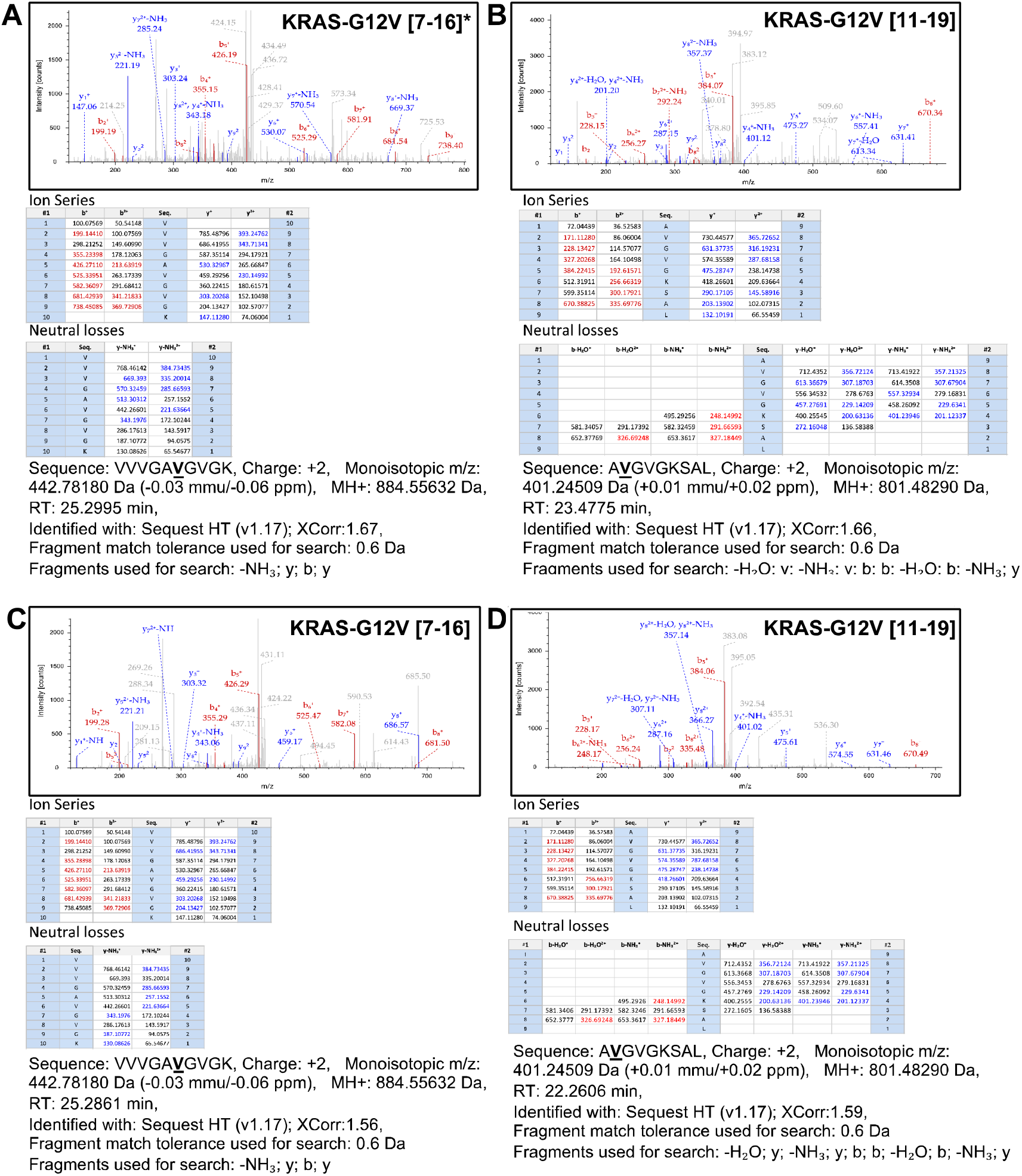
MS2 spectra of targeted-immnopeptidomics for KRAS-G12V. A) The MS2 spectra of oncogenic KRAS-G12V carrying neoantigens, KRAS-G12V [7-16] and B) the neoantigen KRAS-G12V [11-19], in Colo-668. The depicted numbers in the bracket indicates the position of amino acid in KRAS-G12V. The asterisk indicates a positive control sequence previously identified by global-immunopeptidomics analysis. C) and D) The MS2 spectra of oncogenic KRAS-G12V carrying neoantigens, KRAS-G12V [7-16] and KRAS-G12V [11-19], from QGP-1.

**Figure S7.**
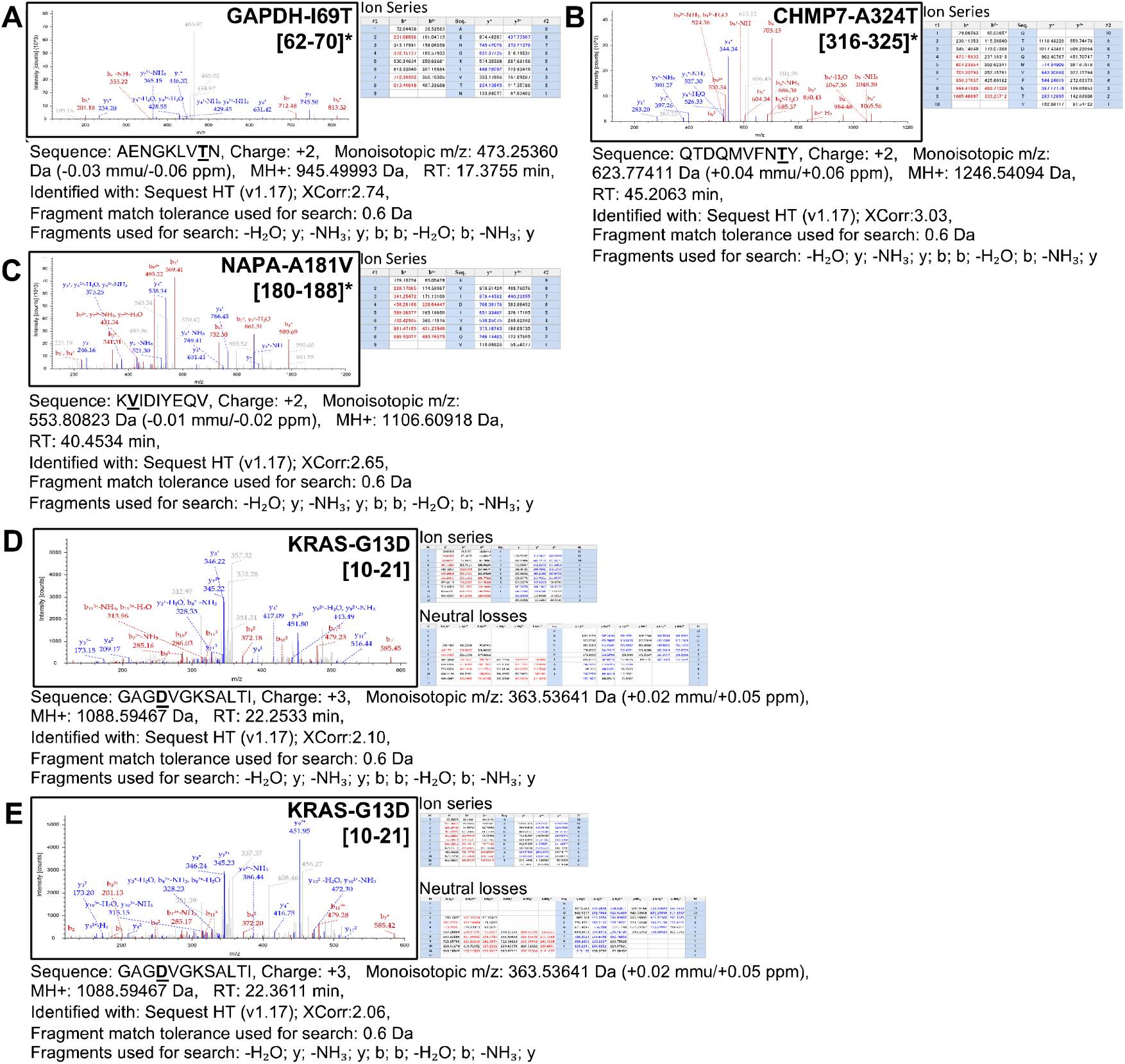
MS2 spectra of newly identified oncogenic KRAS-G13D carrying immunopeptide from HCT116 by targeted-immnopeptidomics. A) to C). MS2 spectra of positive control sequences from neoantigens identified by global-immunopeptidomics from previous studies. The numbers in parentheses indicate the positions of amino acids in the source protein, respectively. The asterisk indicates a positive control sequence previously identified by global-immunopeptidomics analysis. D) and E) Representative MS2 spectra of oncogenic KRAS-G13D carrying neoantigens identified from HCT116 by targeted-immunopeptidomics analysis.

